# Combinatorial Screen of Dynamic Mechanical Stimuli for Predictive Control of MSC Mechano-Responsiveness

**DOI:** 10.1101/2020.12.07.414839

**Authors:** Haijiao Liu, Jenna F. Usprech, Prabu Karthick Parameshwar, Yu Sun, Craig A. Simmons

## Abstract

Mechanobiological-based control of mesenchymal stromal cells (MSCs) to aid in the engineering and regeneration of load-bearing tissues requires systematic investigations of specific dynamic mechanical stimulation protocols. Using deformable membrane microdevice arrays paired with combinatorial experimental design and modeling, we systematically probed the individual and integrative effects of mechanical stimulation parameters (strain magnitude (STRAIN), rate at which strain is changed (RATE) and duty period (DUTY)) on myofibrogenesis and matrix production of MSCs in 3D hydrogels. These functions were found to be dominantly influenced by a novel and higher-order interactive effect between STRAIN and DUTY. Empirical models based on our combinatorial cue-response data predicted an optimal loading regime in which STRAIN and DUTY were increased synchronously over time, which was validated to most effectively promote MSC matrix production. These findings inform the design of loading regimes for MSC-based engineered tissues and validate a broadly applicable approach to probe multifactorial regulating effects of microenvironmental and mechanobiological cues.

## Introduction

Mechanical stimulation potently promotes the growth and maturation of tissues [1][2] and is frequently applied when engineering load-bearing tissues to achieve functional mechanical properties [3]–[5]. In many native and engineered connective and cardiovascular tissues, mechanically-stimulated tissue production results from the mechano-responsiveness of mesenchymal cells, including multipotent mesenchymal stromal cells as a source for engineered tissues [1], [6]–[8]. Improved understanding of the effects of dynamic mechanical stimulation on the regulation of MSC fate and functions is important for effective functional tissue regeneration [9]–[11].

To date, mechanobiological-based control of MSCs to, for example, maximize matrix production towards functional tissues [2], has been largely based on best guesses and one-factor-at-a-time approaches, without systematic investigation of the effect of complex dynamic mechanical stimulation protocols on mechanobiological responses of MSCs [12][13]. In particular, although mechanical stimulation parameters like strain magnitude (STRAIN) [12], the rate of strain change (RATE) [3], and duty period (DUTY) [14][15] have been explored individually, their interactions and integrative effects on MSC fate and functions, as required to generate guidelines applicable to engineered tissues, have not [1], [3], [16].

Biomaterial array platforms enable systematic investigations of complex cue-response networks with multivariate control of the cellular microenvironment [17]–[20]. However, few biomaterial array platforms have integrated mechanical stimulation for mechanobiology studies, despite the critical regulatory effects of mechanical stimulation on cell and tissue functions [6]. To address this limitation, we and others have previously developed deformable membrane array platforms to study the mechanobiological responses of cells to combinations of environmental cues in 2D [21], [22]. The deformable membrane platforms developed by us and stretchable substrate array platforms developed by others have also been adapted to enable 3D mechanical stimulation of cell-seeded biomaterial constructs [23]–[25].

Here we use microdevice arrays and parametric modeling with combinatorial experimental design to identify specific combinations of mechanical loading factors that best promote myofibrogenesis and collagen production of MSCs. We report that MSC mechano-responsiveness is dominated by a novel interaction between strain magnitude and duty period (STRAIN*DUTY). Empirical models based on our combinatorial cue-response data predicted an unexpected optimal loading regime in which STRAIN and DUTY were increased synchronously over time, which was confirmed to maximally promote collagen production by MSCs. Thus, this unique combinatorial screening approach generated novel insights otherwise not available, leading to the identification of a novel mechanical loading regiment for MSCs-based tissue engineering. We expect this approach will be broadly applicable to systematically identify combinations of other mechanobiological cues that optimally guide cell functions in context-specific niches.

## Results

### 1. Combinatorial mechanical stimulation of cell-laden hydrogel constructs

Deformable membrane devices were fabricated to house arrays of optically patterned and covalently bound poly(ethylene glycol)-norbornene (PEG-NB) hydrogels (Figure 1A and S1A). Permutations of strain change rate (RATE or R in condition acronyms), initial strain magnitude (STRAIN or ℇ), and duty period (DUTY or D) were imposed to the MSC-laden gels throughout the culture period (Figure 1B, Table S1). Covalent bonding between the gels and substrate membrane transmitted the cyclic deformation and generated up to 16% nominal tensile strain in the gels (Figure 1C, Movie S1). Fluorescent images of cells were acquired and reconstructed into 3D surfaces for the analysis of cell differentiation and matrix production (Figure 1D). The fluorescent surfaces were batch processed for defined metrics which were then modeled using least squares estimation and parametric regression techniques (Table S2).

**Figure 1.**
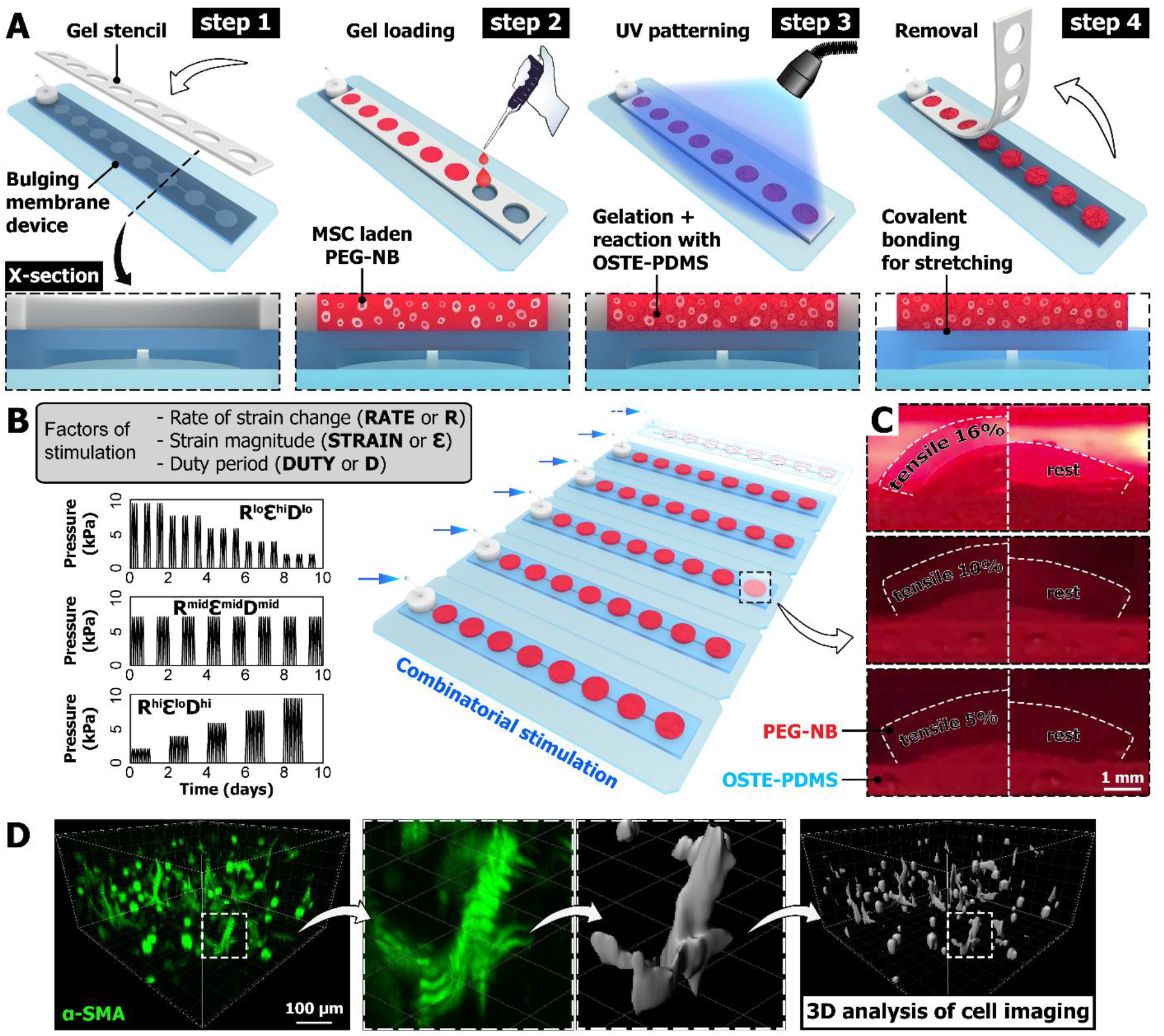
Combinatorial mechanical stimulation of cell-laden hydrogel constructs. (A) Engineered hydrogel construct arrays are optically patterned and covalently bound to microdevice arrays with deformable membranes, to allow for 3D mechanical stretching of cells seeded in the gels. (B) Examples of actuation pressure profile to achieve combinatorial mechanical stimulation patterns in combination of defined levels of RATE,STRAIN,DUTY or R,ℇ,D. (C) Side-view of PEG-NB gels in culture deformed by the membrane under the prescribed actuation pressure, achieving 16%, 10% and 5% of nominal tensile strain. For comparison, the right half image of each group (white dashed line) shows the same gel at rest. (D) For each laser channel (e.g. SMA in green), 3D surfaces are generated using 3D image analysis and automatic thresholding in Imaris (gray). The parametric analysis on various metrics from the fluorescent images and surfaces are performed to assess mechanobiological responses of MSCs (e.g., myofibroblast differentiation and matrix production).

### 2. Interaction between STRAIN and DUTY dominated MSC mechano-responsiveness

As a model, we studied mechanically-stimulated matrix production and myofibroblast differentiation, since myofibroblasts are the critical effectors of tissue remodeling during normal development, repair, and fibrotic conditions [26]–[28]. Since the formation of α-SMA stress fiber is the hallmark of myofibroblast differentiation (Figure 2A, 2I and S1B) [29], we first analyzed the proportion of α-smooth muscle actin stress fiber positive cells (SMA+ proportion) in response to individual and interactive effects of strain change rate, magnitude, and duty period over one week of mechanical stimulation. While the individual factors each had significant effects, parametric analysis on SMA+ cells revealed the STRAIN*DUTY interaction had the largest effect (Figure 2B, Table S2), indicating that combining STRAIN and DUTY both at low or high levels (i.e., positive interactions in patterns (R^hi^ℇ^lo^D^lo^) and (R^hi^ℇ^hi^D^hi^)) led to a high proportion of SMA+ cells (red in Figure 2C; Figure 2I). In contrast, mismatched levels of STRAIN and DUTY (e.g., pattern (R^lo^ℇ^lo^D^hi^) in Figure 2I) or their negative interaction led to low SMA+ proportion (blue in Figure 2C). Mechanical stimulation also increased spreading in MSCs (Figure 2I and S1B) and increased the expression of collagen at local cell protrusions and edges (rim of cell in Figure 2D and 2I zoom-in) compared to those in static culture. Cells that stained for collagen were defined as ‘collagen positive’ (Col+). The parametric analysis on Col+ proportion and collagen intensity consistently revealed that the STRAIN*DUTY interaction as the most sizeable effect (Figure 2E and 2G). Combining STRAIN and DUTY both at low or high levels, i.e., positive STRAIN*DUTY (e.g., patterns (R^hi^ℇ^lo^D^lo^) and (R^hi^ℇ^hi^D^hi^)), led to high Col+ proportion and collagen intensity (red in Figure 2F and 2H), while mismatched levels of STRAIN and DUTY, i.e., negative STRAIN*DUTY (e.g., patterns (R^lo^ℇ^lo^D^hi^) and (R^lo^ℇ^hi^D^lo^)), led to low collagen production (blue in Figure 2F and 2H).

**Figure 2.**
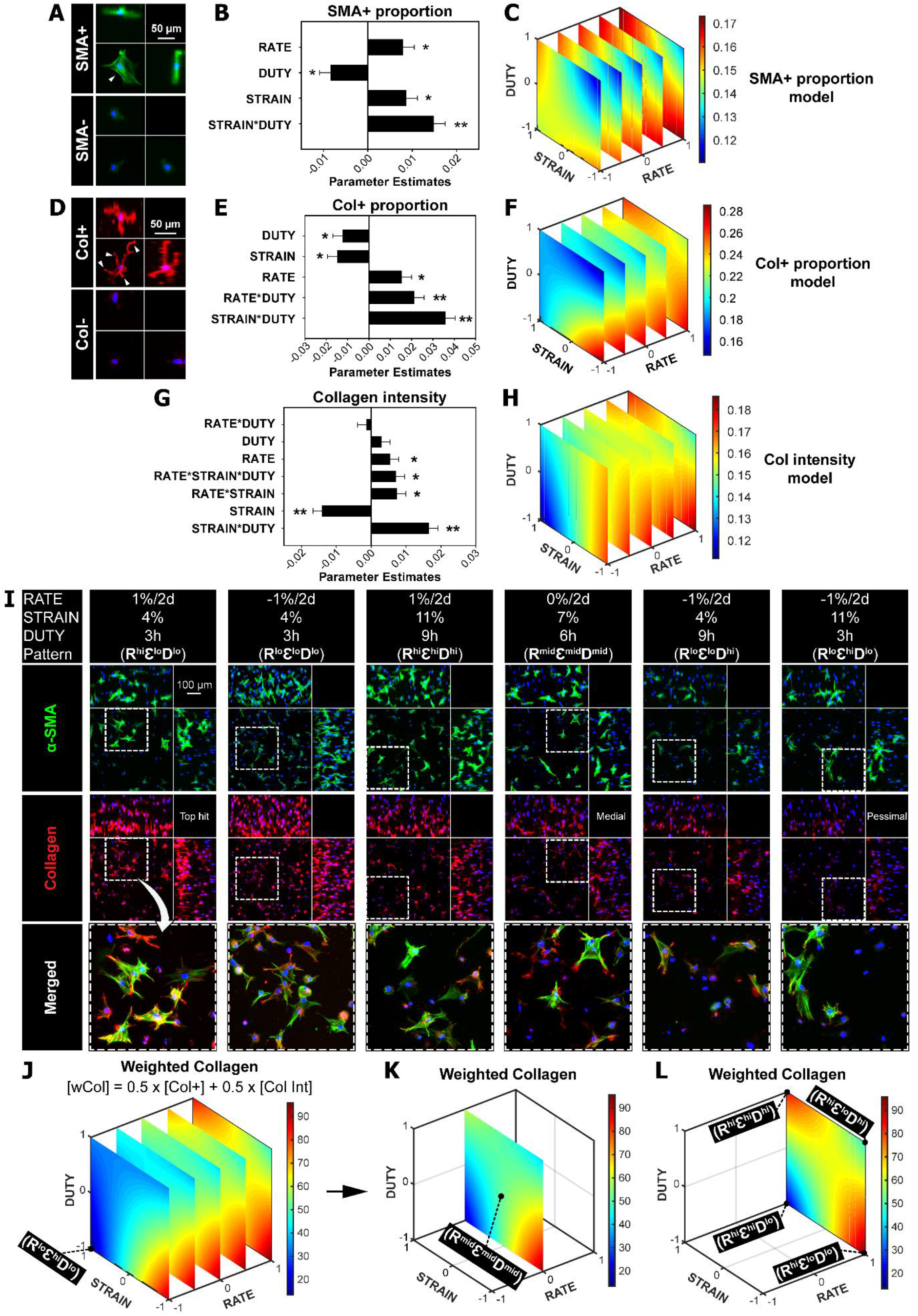
Interaction between STRAIN and DUTY dominated MSC mechano-responsiveness. Parametric models of MSC responses for (A-C) proportion of SMA+ cells, (D-F) proportion of Col+ cells, and (G-H) collagen mean intensity after one-week full factorial screening. (A) Example of a SMA+ cell with visible α-SMA stress fiber formation (top)(arrow head) and a SMA-cell (bottom). SMA in green and nucleus in blue. (D) Example of a Col+ cell with visible collagen (in red) at the spreading cell protrusions and edges (top)(arrow heads) and a Col-cell (bottom). (B, E and G) Summaries of factor effects (parameter estimates) for proportion of SMA+ cells, proportion of Col+ cells and collagen mean intensity. Larger coefficient magnitude of the parameter estimate corresponds to a larger effect of that factor on the model output. (C, F and H) 4D response surfaces plot of the interaction between RATE, STRAIN and DUTY on SMA+ proportion, Col+ proportion and collagen mean intensity. The definition of the coded values is provided in Table S1. (I) Representative maximum intensity projections of cells stained with α-SMA and collagen type I among various patterns of condition. Peripheral images are side view projections. Refer to Table S2 for the polynomial models of SMA+ proportion, Col+ proportion and collagen mean intensity. Error bars represent standard error of the estimated parameters. N = 3-4 independent gels per condition. **p < 0.0001, *p < 0.05. (J-L) The new metric Weighted Collagen (wCol) was defined to identify the top hit, medial and pessimal conditions for further validation. Conditions selected are labeled on the graph and listed in Table S3. (J) 4D response surfaces plot of the interaction between RATE, STRAIN and DUTY on wCol. (K-L) Response surfaces of wCol with RATE at 0%/2d (K) and at 1%/2d (L).

### 3. Harnessing the effect of STRAIN*DUTY is hypothesized to enhance matrix production by MSCs

#### 3.1. Hit conditions were selected for longer term validation

For engineered load-bearing tissues, mechanical stimulation is typically applied more than one week to promote tissue growth. Therefore, we sought to confirm whether selected hit conditions for collagen production from one week culture could maintain their longer term effects. To simplify the assessment of collagen production, a metric ‘Weighted Collagen’ (wCol) was defined based on equal weighting of proportion of Col+ cells and collagen intensity (see Methods section). All conditions were evaluated for levels of wCol to identify the optimal, medial and pessimal conditions of collagen production (Figure 2J). Patterns (R^hi^ℇ^lo^D^lo^), (R^mid^ℇ^mid^D^mid^) and (R^lo^ℇ^hi^D^lo^) that respectively produced the greatest wCol (95/100, Figure 2I and 2L), the medial level wCol (50/100, Figure 2I and 2K) and the lowest wCol (13/100, Figure 2I and 2J) were selected for further validation with extended two-week culture. Notably, the individual effect of strain change rate (RATE) and the interactive effect of STRAIN*DUTY were both significant and positive for all metrics evaluated (Figure 2, S2). In particular, the STRAIN*DUTY interaction had its maximal effect for both SMA and collagen expression at high RATE of 1%/2d (diagonal distribution of red in Figure 2C, 2F, 2H and 2L). This is because the positive effects of RATE and STRAIN*DUTY were additive. In contrast, the effects of other factors such as STRAIN and RATE*DUTY were mostly inconsistent and/or insignificant. Therefore, additional conditions at high RATE of 1%/2d (i.e., patterns (R^hi^ℇ^hi^D^hi^), (R^hi^ℇ^lo^D^hi^) and (R^hi^ℇ^hi^D^lo^)) were selected and added to pattern (R^hi^ℇ^lo^D^lo^) to form a full factorial design to validate the above identified effect of STRAIN*DUTY interaction on MSC responses (Figure 2L). Final conditions selected for the two-week validation culture are listed in Table S3.

#### 3.2. STRAIN*DUTY dominantly affects myofibroblast differentiation and matrix production after two weeks of culture

Prediction expressions of the parametric models for each metric analyzed are shown in Table S4. Parametric analysis of SMA+ proportion revealed that by two weeks (Figure 3A and 3B) the STRAIN*DUTY interaction was still positive and had the greatest effect (Figure 3C; Table S4), generating a saddle-shaped response surface (Figure 3D). The SMA+ proportion increased with STRAIN and DUTY both at high or low levels (red in Figure 3D) and peaked at 19.1% with RATE, STRAIN and DUTY at 1%/2d, 4%, and 3h ON/OFF (i.e., pattern (R^hi^ℇ^lo^D^lo^) in Figure 3B and S4). To complement SMA+ as a marker of myofibroblasts, we also analyzed the expression of fibronectin as an early indicator of activation of myofibroblasts [30]–[32]. Similar to SMA+, fibronectin intensity increased with positive STRAIN*DUTY interaction (Figure 3E, red in Figure 3F, pattern (R^hi^ℇ^hi^D^hi^) in Figure 3B, S4 and S5B). In contrast, mismatched levels of STRAIN and DUTY inhibited both SMA+ proportion and fibronectin intensity (blue in Figure 3D and 3F, pattern (R^hi^ℇ^hi^D^lo^) in Figure 3B and S4). For collagen production, the STRAIN*DUTY interaction remained a positive and dominating effect after parametric analysis of all collagen metrics including Col+ proportion, collagen intensity, normalized volume and integrated density (Figure 3G, 3I, S6A and S6F; Table S4). Collagen production, measured by all the collagen metrics, peaked at high levels of STRAIN and DUTY and plunged with mismatched levels of STRAIN and DUTY (i.e., pattern (R^hi^ℇ^hi^D^hi^) vs. (R^hi^ℇ^hi^D^lo^) in Figure 3B and S4; red vs. blue in Figure 3H and 3J; Figure S5 and S6). Interestingly, evaluation of collagen production by wCol identified the pattern (R^hi^ℇ^hi^D^hi^) that replaced the pattern (R^hi^ℇ^lo^D^lo^) as the new top hit condition by two weeks (wCol = 100). This was due to more significant increases from one week to two weeks in normalized Col+ proportion and collagen intensity of pattern (R^hi^ℇ^hi^D^hi^) (1.9 to 9.3-fold and 1.2 to 2-fold, respectively) than those of pattern (R^hi^ℇ^lo^D^lo^) (2.2 to 6.4-fold and 1.3 to 1.6-fold, respectively) (Figure S7B-C). Additionally, the pattern (R^hi^ℇ^hi^D^lo^) replaced the pattern (R^lo^ℇ^hi^D^lo^) as the new pessimal condition (wCol = 0), which showed little effect at promoting collagen production by two weeks (Figure S7B-C).

**Figure 3.**
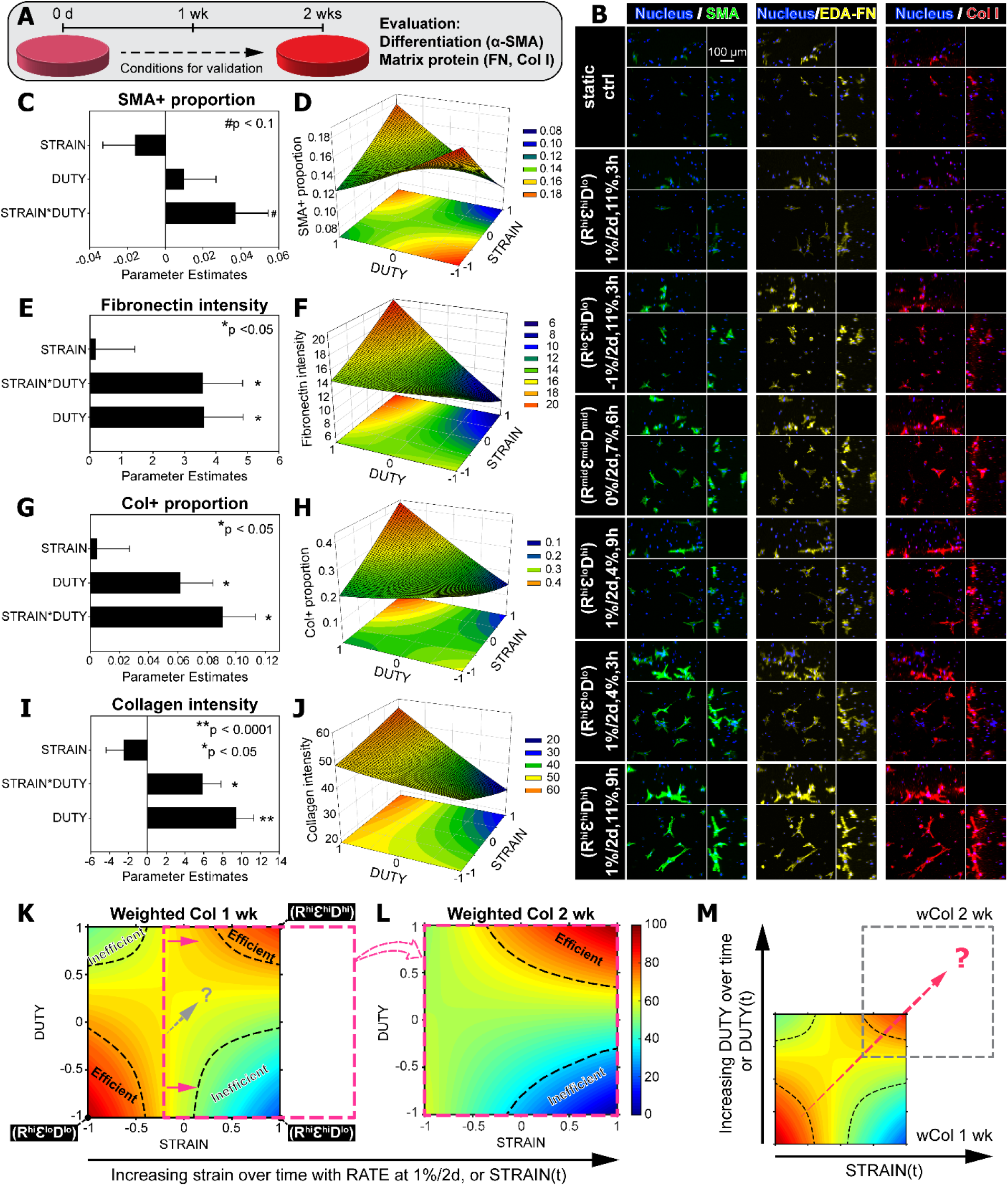
Empirically-driven model generated a hypothesis to enhance matrix production. (A) Overview of the two-week validation culture of selected conditions. (B) Representative maximum intensity projections of MSCs stained with α-SMA, EDA-FN, and collagen type I. Peripheral images are side view projections. (C-J) Parametric analysis of MSC responses in expression of (C-D) α-SMA, (E-F) fibronectin, and (G-J) collagen with RATE at 1%/2days. (C, E, G and I) Summaries of factor effects (parameter estimation) for SMA+ cell proportion, fibronectin intensity, Col+ cell proportion, and collagen intensity, respectively. Error bars represent standard error of the estimated parameters. N = 3-4 independent gels per condition. **p < 0.0001, *p < 0.05, #p < 0.1. (D, F, H and J) Response surfaces of the STRAIN*DUTY interaction on SMA+ proportion, fibronectin intensity, Col+ proportion, and collagen intensity, respectively. 2D projections of the response surfaces are shown at the bottom in each plot. (K) Response surface of Weighted Collagen by one week with highlighted regions of efficient stimulation and inefficient stimulation (dashed outline)(RATE at 1%/2days). (L) Response surface of wCol after two weeks validation culture with regions of efficient and inefficient stimulation, shifted from that of one week in the direction of increasing STRAIN (pink arrows and outline). (M) Differences in response surfaces of wCol between one-week and two-week, due to dynamic changes in the STRAIN*DUTY interaction effect, generated a hypothesis to optimally promote collagen production by MSCs (dashed arrow).

#### 3.3. Increasing STRAIN led to dynamic changes of STRAIN*DUTY interaction

The above parametric analyses revealed that STRAIN*DUTY interaction but not individual STRAIN or DUTY was the dominant determinant of MSC mechanobiological responses including the expression of α-SMA, fibronectin and collagen. Interestingly, comparison of the wCol response surfaces at high RATE of 1%/2d from the initial one week screening (Figure 3K) and the two weeks validation (Figure 3L) showed significant differences. We categorized conditions with wCol > 75 as ‘Efficient’ to mark their promotive effect on high levels of collagen production, and conditions with wCol < 50 as ‘Inefficient’ due to the little effect of mismatched levels of STRAIN and DUTY. After two weeks, patterns (R^hi^ℇ^hi^D^hi^) (i.e., 1%/2d, 11%, 9h ON/OFF) and (R^hi^ℇ^hi^D^lo^) (i.e., 1%/2d, 11%, 3h ON/OFF) remained in their respective categories of ‘Efficient’ and ‘Inefficient’ (Figure 3K and 3L), resulting in the new top hit and pessimal conditions. In comparison, the initial hit condition pattern (R^hi^ℇ^lo^D^lo^) shifted out of its category of ‘Efficient’ with the new wCol only at medial levels (pink arrows and outline in Figure 3K). This was due to the increasing strain magnitude (when RATE at 1%/2d) from the initial 4% to 10% after two weeks matching the unchanged duty period, thus causing dynamic changes in the effect of STRAIN*DUTY interaction from positive to negative (i.e., initially (R^hi^ℇ^lo^D^lo^) to effectively (R^hi^ℇ^hi^D^lo^) by two weeks). Therefore, a reasonable deduction was that the initial hit condition (R^hi^ℇ^lo^D^lo^) may outperform the new top hit condition (R^hi^ℇ^hi^D^hi^) by two weeks, provided the negative change of STRAIN*DUTY interaction can be mitigated. Collectively, this led us to hypothesize that increasing duty period to synchronize with increasing strain magnitude would yield a new condition that maintains the positive effect of STRAIN*DUTY interaction and consistently and efficiently promotes collagen production (dashed arrow in Figure 3K and 3M).

### 4. Synchronous application of STRAIN and DUTY promoted collagen production

To test our hypothesis, we repeated the two-week culture and added a new condition pattern (R^hi^ℇ^lo^D^lo^ sync) that matched the increasing STRAIN to a stepwise increasing DUTY (i.e., 3, 6 and 9 hrs ON/OFF at day 2, 6 and 12, respectively). Compared to the original pattern (R^hi^ℇ^lo^D^lo^) (Figure 4A left), the new pattern (R^hi^ℇ^lo^D^lo^ sync) by design changed the STRAIN, DUTY from initially 4%, 3h to 7%, 6h after one week and to 10%, 9h after two weeks (Figure 4A right) thus maintaining the positive effect of STRAIN*DUTY. By analyzing both collagen and fibronectin, the results confirmed that the new pattern (R^hi^ℇ^lo^D^lo^ sync) indeed outperformed the previous conditions at promoting matrix production. Specifically, the pattern (R^hi^ℇ^lo^D^lo^ sync) significantly and maximally promoted Col+ proportion and collagen intensity to 47.7% ± 1% and 24.2% ± 5% respectively, compared to the previous hit conditions in the initial screen (i.e., pattern (R^hi^ℇ^lo^D^lo^)) and in the validation culture (i.e., pattern (R^hi^ℇ^hi^D^hi^)) (Figure 4B, 4C and 4F top). The corresponding wCol was also significantly higher for pattern (R^hi^ℇ^lo^D^lo^ sync) than for pattern (R^hi^ℇ^hi^D^hi^) and (R^hi^ℇ^lo^D^lo^) (Figure 4D and 4F top). Additional analysis of fibronectin also showed that the pattern (R^hi^ℇ^lo^D^lo^ sync) significantly promoted fibronectin intensity by 60% compared to pattern (R^hi^ℇ^lo^D^lo^), with marginal promotion compared to pattern (R^hi^ℇ^hi^D^hi^) (Figure 4E and 4F bottom).

**Figure 4.**
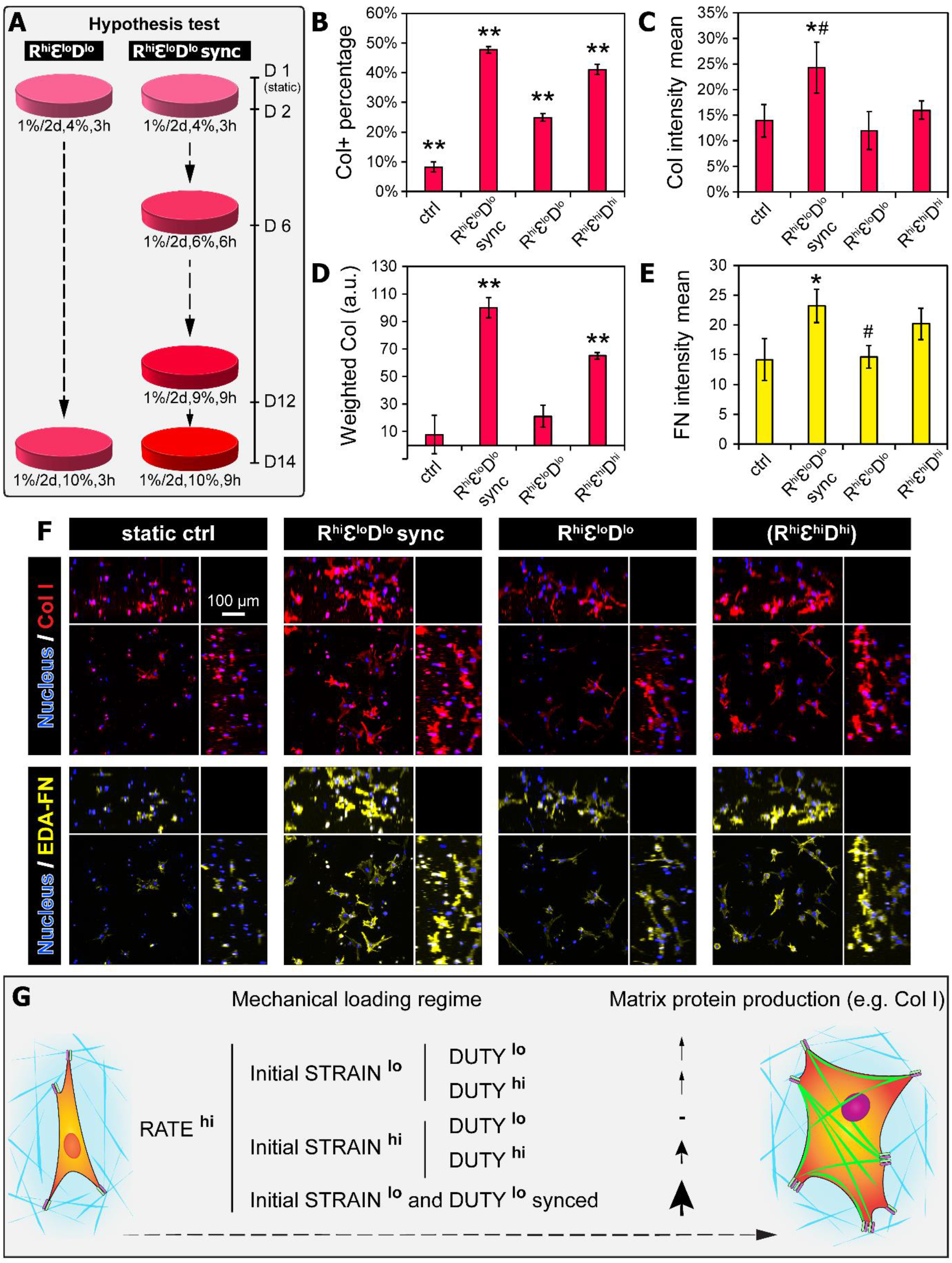
Synchronous application of STRAIN and DUTY promoted collagen production. (A) Overview of the hypothesis test on the new condition for collagen and fibronectin production by MSCs. (left) the original pattern (R^hi^Ɛ^lo^D^lo^); (right) the new condition with dynamically synced STRAIN and DUTY - pattern (R^hi^Ɛ^lo^D^lo^ sync). (B-E) Quantification of collagen and fibronectin production using metrics including: (B) Col+ proportion, (C) Collagen intensity, (D) Weighted Collagen, and (E) fibronectin intensity. N = 3-4 independent gels per condition. **p < 0.01 vs. all other groups, *p < 0.05 vs. ctrl and pattern (R^hi^Ɛ^lo^D^lo^), #p < 0.1 vs. pattern (R^hi^Ɛ^hi^D^hi^). (F) Representative maximum intensity projections of cells stained with collagen and ED-A FN from tested conditions. Peripheral images are side view projections. (G) Schematic summarizing the interplay between STRAIN and DUTY, possibly manifested through integrative regulation of force loading rate via molecular clutch and imprinted mechanical memory which ultimately control the mechano-responsiveness of MSCs.

## Discussion

Systematic and combinatorial studies are required for better understanding of mechanobiological cell responses to complex microenvironmental cues in engineered niches. Factorial screening is a common strategy for optimizing context-specific metrics and maximizing statistical power with finite resources. In this work, by using a 3D bioreactor array platform with combinatorial mechanical stimulation factors, we performed full factorial designs to evaluate the main effect of each factor and their higher-order interactions. With parametric modeling we discovered that the interaction effect between strain magnitude and duty period of applied mechanical stimulation (STRAIN*DUTY) dominantly influenced myofibrogenesis and matrix production by MSCs. Furthermore, empirical modeling of higher-order interactions provided insights into the dynamic changes of STRAIN*DUTY interaction. These insights led us to hypothesize a new dynamic mechanical stimulation scheme in which strain magnitude and duty period were synchronously increased throughout the culture period. Indeed, this approach was verified to outperform the original schemes in promoting MSC matrix production.

Our data showed that the individual effect of strain change rate (RATE) was consistently significant and positive. The application of an incremental strain (with positive RATE at 1%/2days) produced more α-SMA, fibronectin and collagen (Figure 2 and S2). This is consistent with previous reports that showed increased collagen content per MSC-like cell in engineered constructs by incremental cyclic mechanical stimulation [3][33]. It was previously shown that the benefit of constant ‘normal’ mechanical stimulation gradually vanishes due to cell adaptation [34]–[36], and the application of incremental strain was believed to mitigate the cell adaptation by “resetting” the mechanosensitivity of cells [37][38]. Building on these earlier studies, our study revealed that the interaction effect between strain magnitude and duty period (STRAIN*DUTY) more dominantly regulated the mechano-responsiveness of MSCs compared to the individual effect of RATE or incremental strain alone (Figure 2). The STRAIN*DUTY interaction can be observed even when RATE was at zero, which is the application of a constant strain stimulation (Figure 2C and 2K). This interaction effect is significant in that it not only reveals novel insights into new “keys” to control the mechano-responsiveness of MSCs for advanced bioprocess control and maturation of engineered tissues but also emphasizes the capability of using our approach to generate novel hypothesis for testing and validation.

The regulation of cell responses to stretching and related force transmission can be explained by the molecular clutch dynamics and the force loading rate, which is a fundamental factor driving the mechanosensitivity of the molecular clutch [39]. Mechanical cues such as substrate rigidity and dynamic substrate stretching can directly regulate the force loading rate and influence several cell fate decisions and functions [39][40]. Increasing strain at a constant stretching frequency (as in this study) or increasing frequency at a constant strain both convert to increasing force loading rate and transmission, similar to that acquired from increasing substrate rigidity. Individual mechanosensors in the clutch model experience these mechanical cues and also determine force transduction for downstream signaling [39]. For instance, various levels of force loading rate induce rapid and/or long-term molecular events such as talin unfolding and integrin-ECM unbinding [16], ATP and other purinergic signaling and gene silencing by chromatin compaction [41]–[43]. Cyclic mechanical stretching and matrix stiffening have also been shown to modulate the activation of multiple transcription factors [40], [44], [45] and promote myofibroblast differentiation of MSC-like cells with SMA expression [21]. Notably, we found significant difference between the expression pattern of SMA and collagen in terms of top hit conditions (Figure 2, 3 and S7). This suggests discrepancy between myofibroblast differentiation of MSCs and their transformation into synthetic phenotype with optimal matrix production, where potential new insights could be generated using similar approach to our study.

Mechanical cues including substrate rigidity and stretching have been shown to induce chromatin remodeling and influence the epigenomics of MSCs [46]–[48] although the specific mechanisms involved remain elusive. Mounting evidence supports the notion that repeated and lasting mechanical stimulation instills memory in stem cells including MSCs. Many mechanical priming-induced cellular events are differentially executed depending on culture history and have been proposed to serve as mechanical memory keepers on different levels, acting together to adjust the mechanosensitivity of MSCs to mechanical perturbation [46], [47], [49]–[51]. For instance, the nuclear translocation of co-transcription factor YAP and chromatin condensation adapt rapidly to mechanical loading but after loading cessation YAP translocates back to the cytoplasm and chromatin remodeling drops to baseline levels, resulting in reversible or weak mechanical memory and nominal changes in gene expression [46], [51]. In comparison, expression level of microRNA miR-21 was robust against acute changes in environmental mechanics due to long half-life of several days, serving as a long-term memory keeper [50]. Importantly, higher magnitude of mechanical cues led to higher magnitude of mechanical memory imprinted in MSCs and more significant extension of the permanency of memory and modulation on cell functions after loading cessation [46], [47], [50]. Similarly, in our study this mechanical memory may be differentially regulated depending on the level of strain magnitude and contribute to the STRAIN*DUTY interaction (Figure 4G). For example, the patterns (R^hi^ℇ^lo^D^hi^) and (R^hi^ℇ^hi^D^lo^) may result in weak and strong memories respectively due to their respective strain levels and force transmission. These memories either do not persist through the high duty period and lead to only intermittent effect (i.e., (R^hi^ℇ^lo^D^hi^)) or contribute to cell adaptation by lowering the mechanosensitivity of MSCs [3], [38] and cause a diminishing effect (i.e., (R^hi^ℇ^hi^D^lo^)), which are both suboptimal. Other patterns with positive STRAIN*DUTY interaction (i.e., (R^hi^ℇ^lo^D^lo^) and (R^hi^ℇ^hi^D^hi^)) seem to balance the mechanical memories and mechanosensitivity of MSCs and result in relatively more significant stimulating effects. Provided the dynamic nature of the imprinted memories due to increasing strain, we were led to hypothesize and demonstrate the strategy of maintaining continuous and accumulative effects of stimulation for matrix production by matching increasing strain with increasing duty period (Figure 4).

The goal of this study was not to reproduce the collagen content and organization of native load-bearing tissues, but to increase the understanding of responses and development of engineered tissue constructs from MSCs to combinatorial and dynamic mechanical stimulation, specifically in terms of myofibrogenesis and matrix production. Myofibrogenesis may be more relevant to the engineering of specific tissues such as the heart valve and to specific disease modeling such as tissue fibrosis. More comprehensive biological analysis can be performed for context specific applications when needed. Note that the strategy of integrating microdevice arrays and parametric modeling to identify the combinatorial and specific effects of mechanical stimulation can be widely applied to other cell sources (e.g., vascular smooth muscle cell) [1], [16]. Other factors of interest can also be included such as fluidic shear stress, scaffold materials and media formulations for combinatorial screening of their high-order effects on engineered (load-bearing) tissue formation and functions. Non-invasive functional analysis on-chip, similar to integrating strain sensors to monitor the stiffness evolution of engineered constructs [52], would be revolutionary at providing new insights into the dynamics of tissue development. Moreover, the PEG-NB hydrogel system permits precise definition of material and biochemical cues [53], and similar combinatorial screening studies can be implemented to understand the specific and integrative effects of critical microenvironmental factors, leading to comprehensive mechanobiology of target cell niches that can serve as a basis for designing next-generation biomaterials.

## Conclusions

This paper reported a combinatorial screening study of the individual and integrative effects of 3D dynamic mechanical stimulation parameters on myofibrogenesis and matrix production by MSCs cultured in 3D hydrogel constructs. Our deformable membrane microdevice arrays were paired effectively with combinatorial experimental design and parametric regression modeling to generate insights otherwise not available. Strikingly, a high-order and significant STRAIN*DUTY interaction effect was discovered that dominantly determined myofibrogenesis and matrix production by MSCs. A novel dynamic loading regime was predicted based on empirically driven models to promote matrix production by MSCs through optimizing their mechano-responsiveness. The optimal regime predicted by our study – pattern (R^hi^ℇ^lo^D^lo^ sync) where strain magnitude and duty period were synchronously increased over time – was validated to most effectively promote collagen and fibronectin production. Our findings significantly inform the design of efficient mechanical loading regimes for MSC-based tissue regeneration. They also represent a clearly generalizable approach to probe multi-factorial regulation of cell fate by combinatorially controlling microenvironmental stimuli.

## Acknowledgements

The AOMF is acknowledged for the use of confocal microscopy for imaging and the Imaris software for data analysis. The authors acknowledge financial support from Canadian Institute of Health Research (CIHR) Operating Grant (MOP-130481), the Ontario Research Fund – Research Excellence program, and Canada Research Chair in Mechanobiology to CAS and Micro and Nanoengineering Systems to YS.

## Materials and Methods

Please refer to SI Materials and Methods for detailed methods and data analysis.

### 1. Device fabrication and integration of poly(ethylene glycol)-norbornene gel array

The deformable membrane microdevice array is based on our previously developed platform [54][52]. Dynamic 3D mechanical stretching was successfully applied to the poly(ethylene glycol)-norbornene (PEG-NB) hydrogel arrays by covalently bonding them to the membrane which are pneumatically actuated. PEG-NB hydrogel system was used for 3D culture of MSCs because its biochemical and material properties are tunable, including adhesion peptide identities and densities, elasticity and degradability [53], [54].

### 2. Human MSC culture

Cryopreserved human bone marrow derived MSCs were obtained from the Texas A&M Health Science Center College of Medicine Institute for Regenerative Medicine at Scott &White through a grant from NCRR of the NIH (Grant # P40RR017447). MSCs at passage 4-5 and complete culture medium containing 81.7% α-MEM with L-glutamine, 16.3% fetal bovine serum, 1% additional L-glutamine and 1% penicillin/streptomycin were used for all experiments.

### 3. Finite element analysis

3D finite element simulations of the bulging membrane-gel system were performed using ANSYS Workbench v14.0 (ANSYS Inc., Canonsburg, PA) to determine the pressure needed for pneumatic actuation to achieve the prescribed strain magnitude.

### 4. Immunofluorescence staining

Myofibroblast differentiation is defined with neo-expression and incorporation of α-smooth muscle actin (α-SMA) into stress fibers [55]. Therefore, single cell immunofluorescence-based analysis is appropriate to identify the proportion of myofibroblasts from a population of cells. Expression of ED-A fibronectin precedes and is necessary for myofibroblast activation, and is suggested to serve as master template for collagen deposition [30], [32], [56]. MSCs embedded in the PEG-NB gels were stained for α-SMA with FITC-conjugated mouse monoclonal anti-human α-SMA (F3777, Sigma; dilution 1:300), cellular extra domain-A (ED-A) fibronectin with mouse monoclonal anti-ED-A fibronectin (IST-9, sc-59826, Santa Cruz Biotech; dilution 1:200), and collagen with rabbit monoclonal anti-collagen type I (ab138492, Abcam; dilution 1:300).

### 5. Confocal microscopy imaging and analysis

A laser scanning confocal microscope (Zeiss LSM710) with a 20× objective (Plan-Apochromat 20×/1.0 DIC, water immersion) was used to acquire optical slices of the 3D hydrogel constructs. Imaris (Bitplane) was used to create automatically thresholded, quantifiable 3D surfaces around stained objects in the hydrogels for each laser channel from confocal image stacks. To simplify the assessment of collagen production, a new metric ‘Weighted Collagen’ was defined based on Col+ proportion and collagen intensity. Each condition was graded and ranked for both Col+ proportion [Col+] and collagen intensity [Col Int], proportionally with highest value to 100 and lowest to 0. The Weighted Collagen [wCol] was defined using an objective function with Col+ proportion and collagen intensity equally weighted to recognize their putatively equal representation of collagen production.

### 6. Factorial design of experiments and statistical analysis

The factorial design of experiments approach was applied to generate insights on the significance of individual factors and interactions between factors (e.g., parameters of dynamic mechanical stimulation – Change rate of strain or RATE, Initial strain magnitude or STRAIN, and Duty period or DUTY) on a given output metric (e.g., α-SMA, collagen or fibronectin expression) [57][20]. A full factorial design included complete combinations of high and low levels of interest for each factor (i.e., RATE from 1%/2days to −1%/2days, STRAIN from 11% to 4%, DUTY with 50% duty cycle from 9hrs ON/9hrs OFF to 3hrs ON/3hrs OFF; Figure 1B; Table S1). Additional combinations involving medial levels of factors or “center points” were included to further derive potential quadratic or curvature effects (i.e., RATE at 0%/2days, STRAIN at 7%, DUTY at 6hrs ON/6hrs OFF; Table S1). The conditions of the three-factor full factorial design were randomized in experiments with at least three replicates per condition. Parametric models were generated on the combinatorial cue-response data and associated statistical analysis were performed all using JMP 13 (SAS Institute). Additional statistical analyses were performed using SigmaPlot 12. Data are reported as mean ± standard error of the mean unless otherwise noted, and were analyzed by one-way and two-way ANOVA with Tukey post hoc tests for all pairwise comparisons and with statistical significance evaluated at p < 0.05.

### 7. Parametric modeling

In a process as described previously [20][58], response surfaces were modeled by linear regression. For example, for a three-factor response surface, the output response (Y) was modeled as a function of independent input factors (X_i_) in a polynomial function Eq. (2),

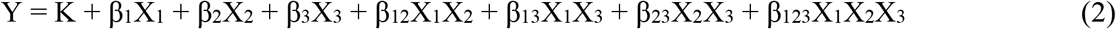

where K corresponds to the average response of center points, β_i_ the main effect coefficients, β_ij_ second-order interaction coefficients. The design matrix and coded values are described in Table S1. Coded values were useful as they allow for the determination of factor effects independent of units through the comparison of β coefficients [57]. Model parameters were estimated using least squares estimation in JMP and the statistical significance of each parameter was evaluated at p < 0.05. Insignificant factors were removed from the initial model by rank and model parameters were calculated for the reduced model through an iterative process of backward elimination [58].

