## Supporting Information for "Combinatorial Screen of Dynamic Mechanical Stimuli for Predictive Control of MSC Mechano-Responsiveness"

### Materials and Methods

#### 1. Device fabrication and integration of poly(ethylene glycol)-norbornene gel array

The bulging membrane device array used is based on our previously developed platform [1][2] that enabled dynamic 3D mechanical stretching of arrays of cell-laden hydrogels. It was applied for the demonstration in culturing 3D cell-laden hydrogel arrays and in combinatorial screening of factors in dynamic 3D mechanical stimulation. Specifically, 3D mechanical stretching was applied to the poly(ethylene glycol)-norbornene hydrogels (PEG-NB) by covalently bonding them to the bulging membranes. We used PEG-NB hydrogel system for 3D culture of MSCs because its biochemical and material properties can be tuned, including adhesion peptide identities and densities, elasticity and degradability [1], [3]. The bulging membrane devices were fabricated from off-stoichiometry thiol-ene based polydimethylsiloxane (OSTE-PDMS) using a composition ratio of 2:0.3:1.5:1.5:1.5 (in grams) plus 0.4 wt% Irgacure TPO-L photoinitiator (BASF). The detailed fabrication process was described previously [1][2]. The model biomaterial PEG-NB was synthesized and crosslinked to bond to OSTE-PDMS as previously described [1]. The cell degradable PEG-NB was decorated with CRGDS adhesion peptides (GenScript) and crosslinked via the matrix metalloproteinase (MMP)-degradable peptide sequence KCGGPQGIWGQGCK, which is flanked with thiol-containing cysteine groups (GenScript), to enable cell adhesion and cell-mediated matrix remodeling. Lyophilized

PEG-NB was dissolved in phosphate buffered saline (PBS) and mixed with the crosslinker, CRGDS peptides and the Irgacure 2959 photoinitiator (BASF) to form the prepolymer solution. Then MSCs suspension were mixed 1:3 with the PEG-NB solution to achieve a final concentration of 11% w/v of PEG-NB, 2.5 mM CRGDS, 0.05% w/v Irgacure 2959, 35% crosslinking density and 1 x $10^6$ cells/mL to form the PEG-NB gels.

To integrate the cell seeded PEG-NB gels onto the OSTE-PDMS bulging membrane device, a thin mortar layer of uncured OSTE-PDMS was applied on top of the device layer first. Then stencils of Sylgard PDMS, containing an array of cylindrical openings (6 mm diameter and 0.7 mm height), were sterilized and aligned on the mortar layer concentrically to the bulging membrane units. Next, 25 µL of the cell-laden PEG-NB solution was added to each stencil well. The cell-laden PEG-NB solution, together with the OSTE-PDMS mortar layer were polymerized under UV light for two minutes. Finally, the rim of cured gels was carefully separated from the stencils wall using tweezers and the stencils were peeled off, leaving the cell seeded PEG-NB hydrogels covalently bound to the OSTE-PDMS membrane via the thiol-ene reaction.

### 2. Human MSC culture

Human MSCs are promising cell sources for regenerative therapies and are used here as a model for mechanosensing studies. Cryopreserved human bone marrow derived MSCs were obtained from the Texas A&M Health Science Center College of Medicine Institute for Regenerative Medicine at Scott &White through a grant from NCRR of the NIH, Grant # P40RR017447. Passage 4-5 MSCs and complete culture medium containing 81.7% α-MEM with L-glutamine, 16.3% fetal bovine serum, 1% additional L-glutamine and 1% penicillin/streptomycin were used for all experiments. MSCs-seeded hydrogel arrays in the device dedicated for 3D mechanical stimulation were cultured statically for one day. Then mechanical stimulation was applied with prescribed nominal tensile strain pattern at 0.1 Hz for 6 days for the initial screening (with strain up to 13%) and for 13 days for the validation experiment (with strain up to 16%). For static conditions, the gels were cultured without mechanical stimulation. Culture media supplemented with 100 µM ascorbic acid was changed every day. The cell-laden hydrogel arrays were cultured in the device in 100 mm Petri dishes and maintained in a humidified 37 ºC incubator with 5% $CO_2$.

### 3. Operation of 3D stretching platform

Diaphragm pumps (Schwarzer, model SP 500EC) and pressure regulators (Marsh Bellofram, model 3410) were integrated into individual sets to pneumatically actuate multiple devices running in parallel to achieve prescribed conditions per device. 3D dynamic mechanical stretching of cell seeded PEG-NB arrays was achieved by applying a triangle waveform pressure with prescribed magnitude at 0.1 Hz. In-house electronics and control scripts were applied to regulate and monitor the cyclic driving pressure, and to achieve the prescribed duty period for each device. To achieve the prescribed change rate of strain magnitude, the regulated pressure magnitude for each device was updated every two days.

### 4. Finite element analysis

3D finite element simulations of the bulging membrane-gel system were performed using ANSYS Workbench v14.0 (ANSYS Inc., Canonsburg, PA) to determine the pressure needed to actuate each device. The PEG-NB gel and OSTE-PDMS were modelled as nearly incompressible isotropic and elastic materials with Poisson's ratio of 0.49. Optical images were experimentally taken using a Navitar zoom system (Navitar, Rochester, NY) and analyzed using ImageJ (NIH), to determine the thickness of OSTE-PDMS device layer and the dimensions of PEG-NB gels as modeling inputs in ANSYS. The OSTE-PDMS membrane typically had a thickness of ~1 mm and a gap distance underneath it of 0.3 mm. The portion of OSTE-PDMS structure without the gap was included beyond the gap region to match the physical structure, similar to the analysis as described previously in [1], [2]. The tensile modulus of OSTE-PDMS (as characterized before in [1]) and elastic modulus of PEG-NB gels (i.e., 48 ± 3.5 kPa) were inputted to the FEA model, and the strain distribution was read as the output of the simulation. To determine the actuation pressure required to achieve various tensile strain levels, the forward FEA based on the material parameter inputs were iterated with a sweep of pressure magnitudes to match the prescribed nominal tensile strain magnitude.

### 5. Immunostaining

To assess the combinatorial effect of 3D mechanical stimulation factors on cell responses, MSCs embedded in the PEG-NB gels were stained for α-smooth muscle actin (α-SMA) as a biomarker for myofibroblasts, cellular extra domain A (ED-A) fibronectin whose expression precedes and is necessary for myofibroblast activation and is suggested to serve as master template for collagen deposition [4]–[6], and collagen type I (Col I) for collagen production. The defining characteristic of differentiated myofibroblasts includes the neo-expression and incorporation of α-SMA into stress fibers [7]. Thus, single cell immunofluorescence-based analysis is appropriate to identify the proportion of myofibroblasts from a population of cells. The PEG-NB gels assigned for immunostaining were washed with TBS and fixed with 10% neutral buffered formalin for one hour at room temperature (RT). The fixed gels were permeabilized with 0.25% Triton X in TBS and blocked with 10% bovine serum albumin for one hour at RT. Then gels were incubated with primary antibodies in TBS with 5% goat serum (GS) overnight at 4 ºC, including rabbit monoclonal anti-collagen type I (ab138492, Abcam; 1:300 dilution) and mouse monoclonal anti-ED-A fibronectin (IST-9, sc-59826, Santa Cruz Biotech; 1:200 dilution). The next day, gels were washed and blocked with 10% GS in TBS for one hour at RT, followed by incubation with secondary antibodies in TBS with 5% GS for 1.5 hours at RT in the dark, including Alexa Fluor 568 goat anti-rabbit (1:300) and Alexa Fluor 647 goat anti-mouse (1:300). Then the gels were washed and incubated with FITC-conjugated mouse monoclonal anti-human α-SMA (F3777, Sigma; 1:300) in TBS with 5% GS overnight at 4 ºC. Next day, gels were washed and incubated with Hoechst 33342 (P162249, Fisher; 1:50) for 15 minutes at RT. Finally, the gels were washed with TBS and deionized water, then covered with Fluoromount for signal preservation prior to imaging with confocal microscopy.

**6. Confocal imaging and analysis**

A laser scanning confocal microscope (Zeiss LSM710) with a 20X objective (Plan-Apochromat 20x/1.0 DIC, water immersion) was used to acquire optical slices of the 3D hydrogel constructs. Each gel was imaged in the middle within a region of 300 μm thickness from the top surface, where the nominal tensile strain was prescribed and evenly distributed [1]. The 3D rendering and analysis software, Imaris (Bitplane), was utilized to create automatically thresholded, quantifiable 3D surfaces around stained objects in the hydrogels for each laser channel from confocal z-stack images. The proportion of cells with visible α-SMA stress fibers formation, or SMA+ proportion, was used to assess myofibroblastic differentiation. The proportion of spreading cells with visible collagen at the cell protrusions and edges, or Col+ proportion, and collagen intensity were both used to assess collagen expression. To simplify the assessment of collagen production, a new metric 'Weighted Collagen' was defined based on Col+ proportion and collagen intensity. Each combination of conditions from the initial screening was graded and ranked for both Col+ proportion [Col+] and collagen intensity [Col

Int], proportionally with highest value to 100 and lowest to 0. The Weighted Collagen [wCol] was defined using an objective function as follows in Eq. (1),

$$[wCol] = 0.5 \times [Col+] + 0.5 \times [Col\ Int] \quad (1)$$

where Col+ proportion and collagen intensity were equally weighted to recognize their putatively equal representation of collagen production. wCol was utilized to select the top hit, medial and pessimal conditions for validation with extended culture of two weeks. In addition, the total volume of collagen, the integrated density of fluorescence intensity of collagen and the fibronectin intensity were also used to evaluate collagen and fibronectin expression. For collagen expression, the total volume of collagen stained surfaces was normalized to the total volume of nuclei in each condition as collagen can be found both intra- and extracellularly. The collagen integrated density of fluorescence intensity was defined as the average product of collagen mean intensity and surface area for all stained surfaces in each image stack. Multiple stacks from each gel were averaged for comparison.

### 7. Factorial design of experiments and statistical analysis

The factorial design of experiments approach was applied to gather information on the significance of both individual factors tested and interactions between factors on a given output (e.g., SMA, collagen or fibronectin) [8][9]. The related data can be used to generate parametric models for statistical analysis and output optimization in a process known as response surface methodology [3][10]. The full factorial design here was used to screen and model the 3D mechanical stretching conditions for their efficacy in inducing matrix production by MSCs in vitro. A full factorial design includes complete combinations of high and low levels for each factor to derive full information on individual and interaction effects of factors. Additional combinations involving a medial value or “center point” can be included to further derive potential quadratic effects (Table S1). A fraction of conditions in the initial screening design matrix were also analyzed for fibronectin expression to constitute a fractional factorial design. The conditions generated in the three-factor full factorial design were randomized in experiments with at least three replicates per condition. Parametric models were then created on the pooled data and relevant statistical analysis were performed using JMP 13 (SAS Institute). All other statistical analyses were performed using SigmaPlot 12. Data are reported as mean ± standard error of the mean (SEM) unless otherwise noted and were analyzed by one-way and two-way

ANOVA with Tukey post hoc tests for all pairwise comparisons. The statistical significance in each comparison was evaluated with $p < 0.05$.

### 8. Parametric modeling

In a process reported previously [9][10], response surfaces were modeled by linear regression using JMP. For example, for a three-factor response surface, the output response (Y) was modeled as a function of independent input factors (Xi) in a polynomial function Eq. (2),

$$Y = K + \beta 1X1 + \beta 2X2 + \beta 3X3 + \beta 12X1X2 + \beta 13X1X3 + \beta 23X2X3 + \beta 123X1X2X3 \quad (2)$$

where K corresponds to the average response of center points, βi the main effect coefficients, and βij the second-order interaction coefficients. The design matrix and coded values are described in Table S1, which represents a three-factor screening experiment with replicates. Coded values were useful as they allow for the determination of factor effects independent of units through the comparison of β coefficients [8]. Model parameters were estimated using least squares estimation in JMP and the statistical significance of each parameter was evaluated with $p < 0.05$. Through an iterative process of backward elimination [10], insignificant factors ($p > 0.05$) were removed from the initial model by rank and model parameters were calculated from the reduced model.

Table S1. Design matrix for the three-factor full factorial design.

| | | **Coded variables** | | | **Actual values** | | |
|---|---|---|---|---|---|---|---|
| Experiment condition | Pattern | RATE or **R** | STRAIN or **$\varepsilon$** | DUTY or **D** | Strain change rate | Initial strain magnitude | Duty period – 50% ON/OFF |
| 1 * | $R^{lo}\varepsilon^{hi}D^{hi}$ | -1 | 1 | 1 | -1%/2days | 11% | 9 hrs |
| 2 | $R^{hi}\varepsilon^{lo}D^{hi}$ | 1 | -1 | 1 | 1%/2days | 4% | 9 hrs |
| 3 * | $R^{mid}\varepsilon^{mid}D^{mid}$ | 0 | 0 | 0 | 0%/2days | 7% | 6 hrs |
| 4 | $R^{hi}\varepsilon^{hi}D^{lo}$ | 1 | 1 | -1 | 1%/2days | 11% | 3 hrs |
| 5 | $R^{lo}\varepsilon^{lo}D^{lo}$ | -1 | -1 | -1 | -1%/2days | 4% | 3 hrs |
| 6 | $R^{mid}\varepsilon^{mid}D^{mid}$ | 0 | 0 | 0 | 0%/2days | 7% | 6 hrs |
| 7 * | $R^{hi}\varepsilon^{lo}D^{lo}$ | 1 | -1 | -1 | 1%/2days | 4% | 3 hrs |
| 8 * | $R^{hi}\varepsilon^{hi}D^{hi}$ | 1 | 1 | 1 | 1%/2days | 11% | 9 hrs |
| 9 | $R^{mid}\varepsilon^{mid}D^{mid}$ | 0 | 0 | 0 | 0%/2days | 7% | 6 hrs |
| 10 * | $R^{lo}\varepsilon^{hi}D^{lo}$ | -1 | 1 | -1 | -1%/2days | 11% | 3 hrs |
| 11 * | $R^{lo}\varepsilon^{lo}D^{hi}$ | -1 | -1 | 1 | -1%/2days | 4% | 9 hrs |

* Conditions marked were additionally analyzed for fibronectin in the initial screening.

Table S2. Prediction expressions for parametric models in the initial one-week screening.

| **Cell responses metric** | **Prediction expression** |
|---|---|
| Col+ cell proportion | $Y = 0.217 + 0.0153 \times Rate + (-0.0146) \times Strain + (-0.0122) \times Duty + 0.0212 \times Rate \times Duty + 0.0357 \times Strain \times Duty$ |
| Col intensity | $Y = 0.152 + 0.00533 \times Rate + (-0.014) \times Strain + 0.00731 \times Rate \times Strain + 0.00286 \times Duty + (-0.00131) \times Rate \times Duty + 0.0165 \times Strain \times Duty + 0.007 \times Rate \times Strain \times Duty$ |
| SMA+ cell proportion | $Y = 0.15 + 0.00787 \times Rate + 0.00859 \times Strain + (-0.00833) \times Duty + 0.0148 \times Strain \times Duty$ |
| FN intensity | $Y = 0.0747 + 0.00686 \times Rate + (-0.00441) \times Strain$ |

Table S3. Conditions selected for the two-week validation culture.

| | | Coded variables | | | Actual values | | |
|---|---|---|---|---|---|---|---|
| Experiment condition | Pattern | RATE or **R** | STRAIN or **Ɛ** | DUTY or **D** | Strain change rate | Initial strain magnitude | Duty period – 50% ON/OFF |
| 2 | $\mathbf{R}^{hi}\mathbf{\varepsilon}^{lo}\mathbf{D}^{hi}$ | 1 | -1 | 1 | 1%/2days | 4% | 9 hrs |
| 3 | $\mathbf{R}^{mid}\mathbf{\varepsilon}^{mid}\mathbf{D}^{mid}$ | 0 | 0 | 0 | 0%/2days | 7% | 6 hrs |
| 4 | $\mathbf{R}^{hi}\mathbf{\varepsilon}^{hi}\mathbf{D}^{lo}$ | 1 | 1 | -1 | 1%/2days | 11% | 3 hrs |
| 7 | $\mathbf{R}^{hi}\mathbf{\varepsilon}^{lo}\mathbf{D}^{lo}$ | 1 | -1 | -1 | 1%/2days | 4% | 3 hrs |
| 8 | $\mathbf{R}^{hi}\mathbf{\varepsilon}^{hi}\mathbf{D}^{hi}$ | 1 | 1 | 1 | 1%/2days | 11% | 9 hrs |
| 10 | $\mathbf{R}^{lo}\mathbf{\varepsilon}^{hi}\mathbf{D}^{lo}$ | -1 | 1 | -1 | -1%/2days | 11% | 3 hrs |

Table S4. Prediction expressions for parametric models in the two-week validation culture.

| **Cell responses metric** | **Prediction expression** |
|---|---|
| Col+ cell proportion | $Y = 0.267 + 0.0045 \times Strain + 0.0617 \times Duty + 0.0906 \times (Strain - 0.167) \times (Duty + 0.167)$ |
| Col intensity | $Y = 44.689 + (-2.445) \times Strain + 9.407 \times Duty + 5.835 \times (Strain - 0.269) \times (Duty + 0.115)$ |
| SMA+ cell proportion | $Y = 0.148 + (-0.0159) \times Strain + 0.00957 \times Duty + 0.0371 \times (Strain - 0.167) \times (Duty + 0.167)$ |
| FN intensity | $Y = 14.428 + 0.165 \times Strain + 3.601 \times Duty + 3.572 \times (Strain - 0.136) \times (Duty - 0.136)$ |
| Col volume | $Y = 2.65 + 0.241 \times Strain + 0.571 \times Duty + 0.809 \times (Strain - 0.25) \times (Duty + 0.25)$ |
| Col integrated density | $Y = 121058.506 + (-655.289) \times Strain + 42821.261 \times Duty + 25966.543 \times (Strain - 0.222) \times (Duty + 0.0741)$ |

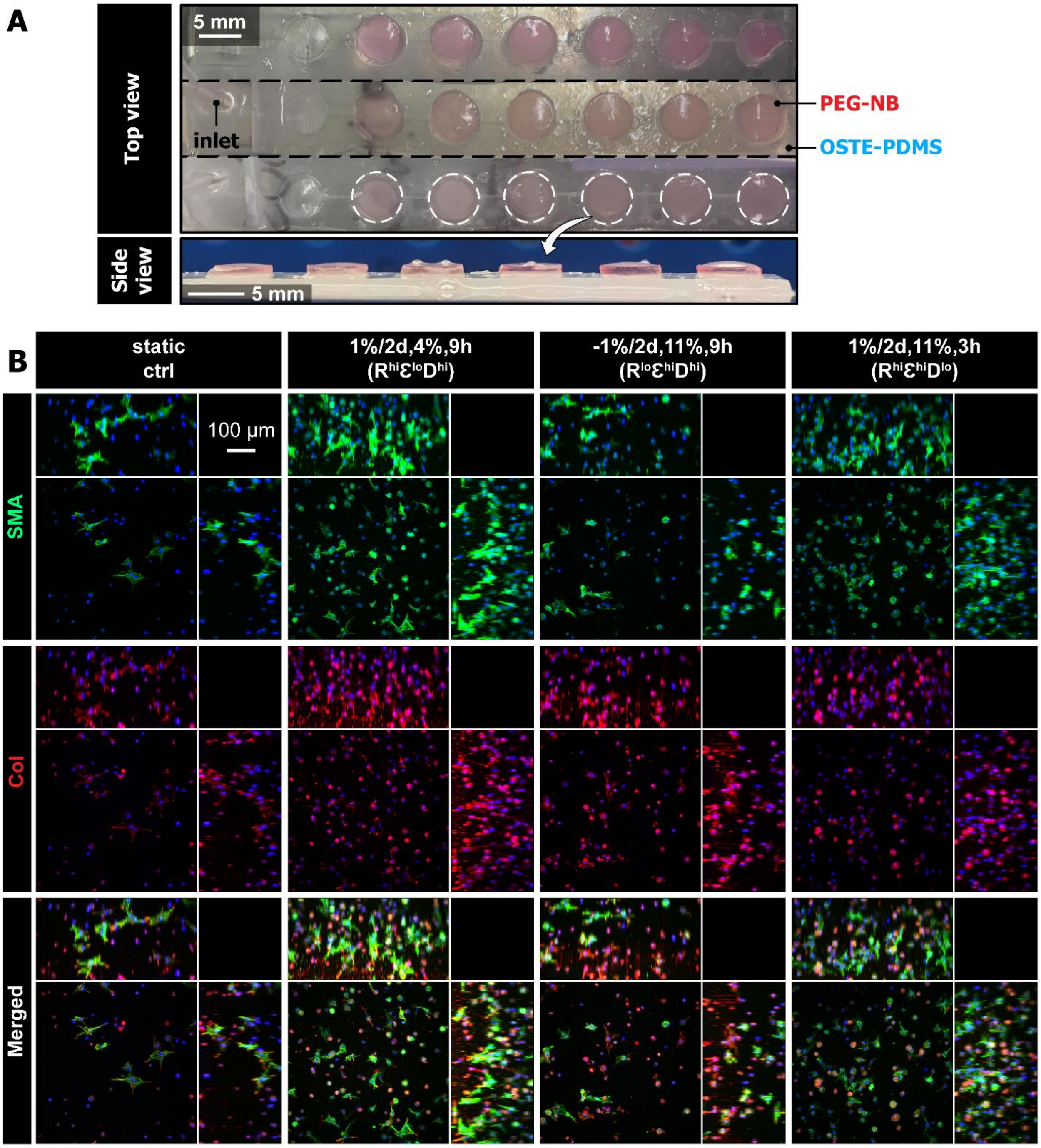


Figure S1. (A) Images of deformable membrane array devices made of OSTE-PDMS with covalently bound PEG-NB gels. (B) MSC responses to additional conditions from the initial one-week factorial screening that are not listed in Figure 2, including patterns ($R^{hi}Ɛ^{lo}D^{hi}$), ($R^{lo}Ɛ^{hi}D^{hi}$), ($R^{hi}Ɛ^{hi}D^{lo}$) and static control. The images show the representative maximum intensity projections of cells stained with α-smooth muscle actin (SMA) and collagen (Col I). Peripheral images are side view projections.

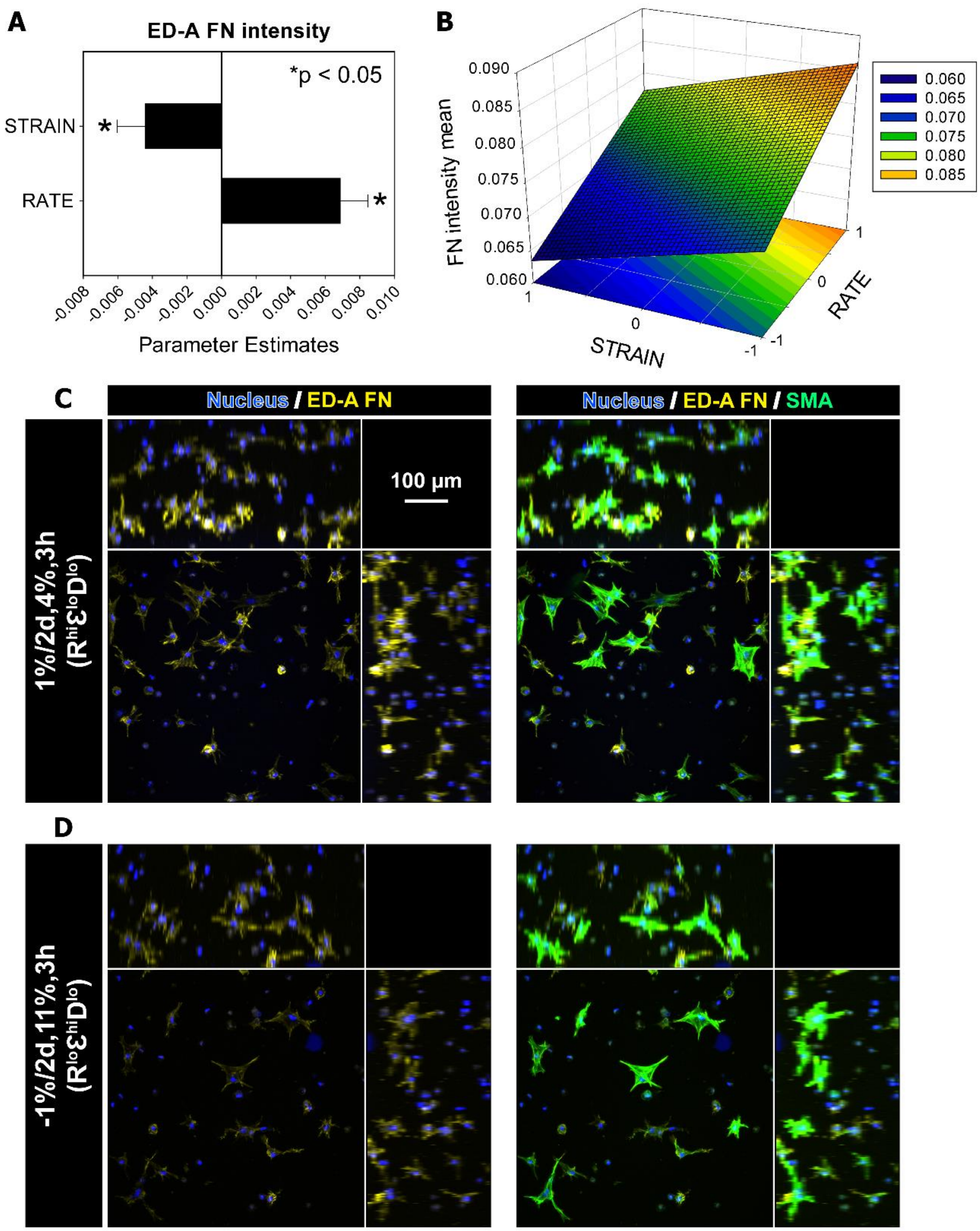


Figure S2. Parametric model of MSC responses of fibronectin production to dynamic mechanical stimulation from the one-week fractional factorial screening (i.e., marked conditions in Table S1). (A) A summary of parameter estimates for FN intensity. Refer to Table S2 for the prediction expression for the response of FN intensity. N = 3-4 replicated gels per condition. (B) Response surface plot on FN intensity as a function of RATE and STRAIN. 2D projection of the response surface is shown at the bottom of plot. The definition of the coded values is provided in Table S1. (C-D) Representative maximum intensity projections of cells stained with ED-A FN and α-SMA. (C) Pattern ($R^{hi}\varepsilon^{lo}D^{lo}$) with high FN intensity and (D) Pattern ($R^{lo}\varepsilon^{hi}D^{lo}$) with low FN intensity. Peripheral images are side view projections.

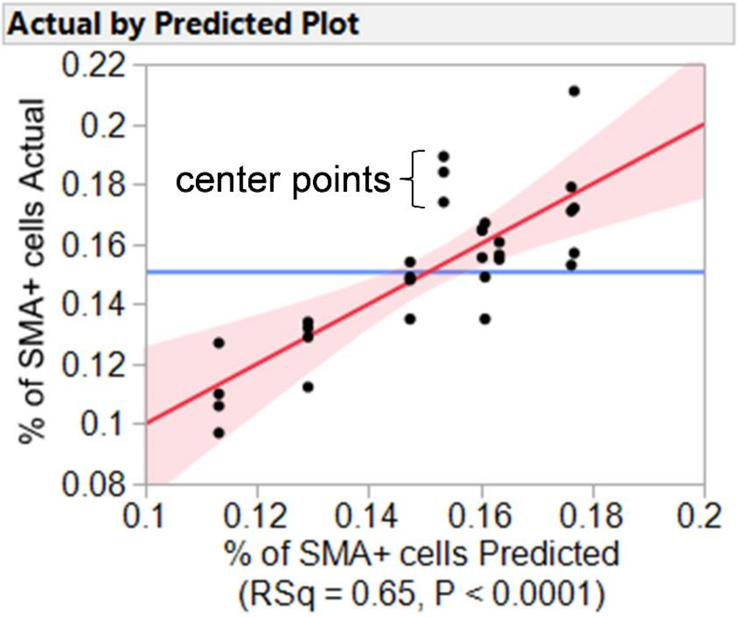


Figure S3. The plot of SMA+ cell proportion by one-week, comparing the actual values (y axis) for each condition or pattern (i.e., each column of dots) with the model predicted values (x axis). The center points ($R^{mid}\varepsilon^{mid}D^{mid}$) did not fit into the linear correlation between the SMA+ proportion and the significant main effects (red line) identified in Figure 2B, which suggested a curvature effect within the levels of factor tested here for RATE, STRAIN and DUTY. Each dot represented one data point averaged from one biological replicate of gel. The red line indicated the line of fit with the filled region in pink indicating p = 0.05 significance curve. The blue line indicated the mean of all data.

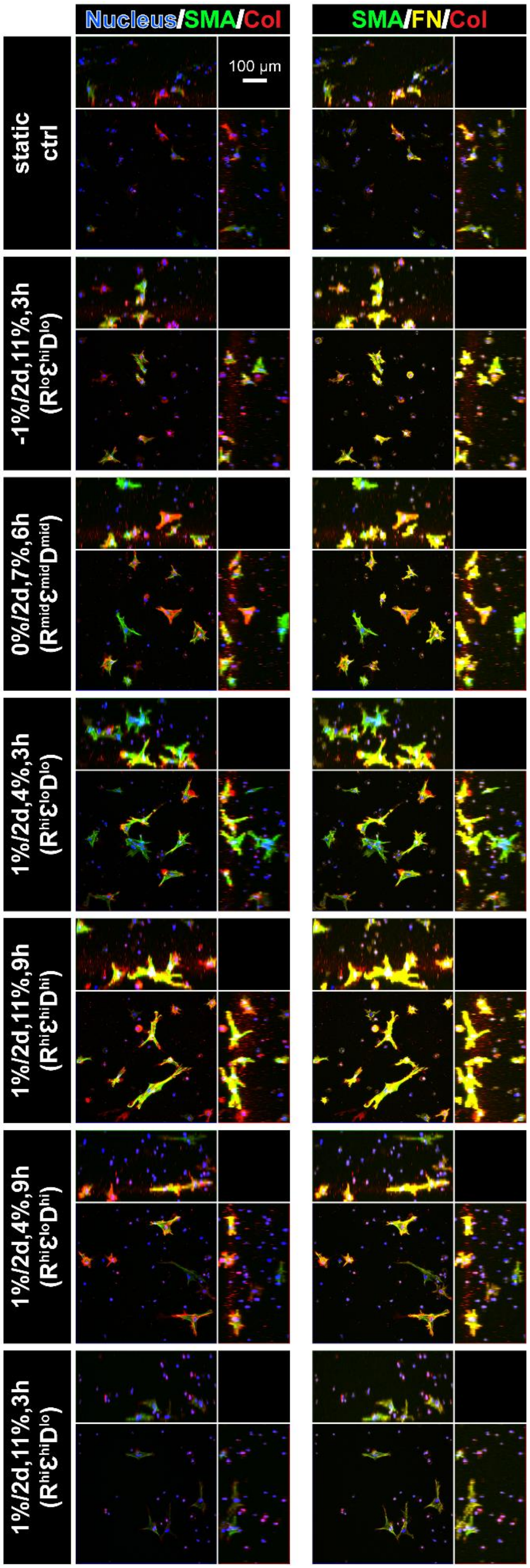


Figure S4. Multi-channel merged views of MSC responses to selected conditions from two-week validation culture. Representative maximum intensity projections of cells stained with α-smooth muscle actin (SMA), collagen (Col I), and fibronectin (ED-A FN). Peripheral images are side view projections.

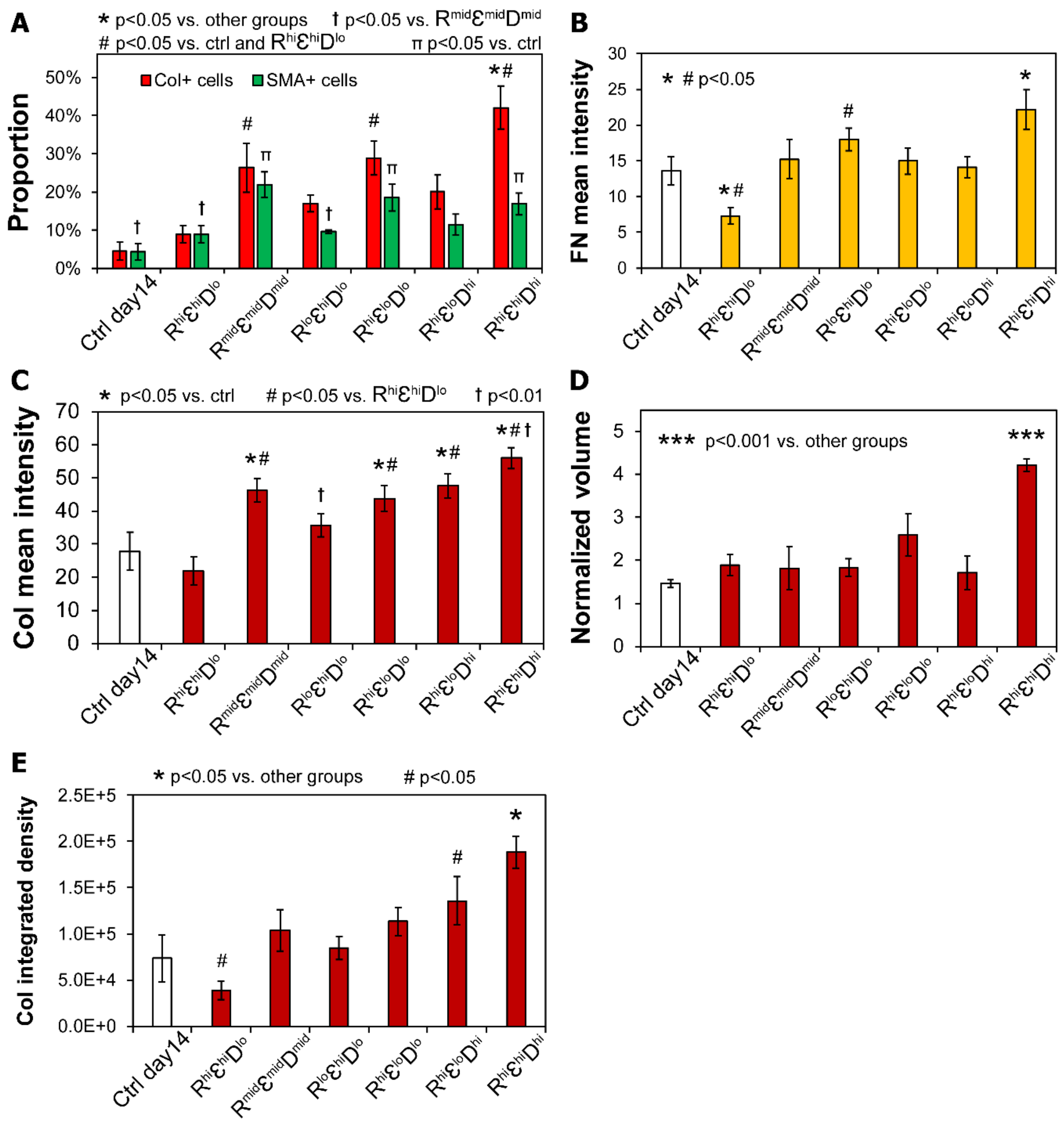


Figure S5. Quantification and pairwise comparison of MSC responses to selected conditions from two-week validation culture. (A) Col+ and SMA+ cells proportion. *p<0.05 vs. other groups, #p<0.05 vs. ctrl and pattern ($R^{hi}\varepsilon^{hi}D^{lo}$), † p<0.05 vs. pattern ($R^{mid}\varepsilon^{mid}D^{mid}$), π p<0.05 vs. ctrl. (B) FN mean intensity. *p<0.05 and #p<0.05 between marked groups. (C) Collagen mean intensity. *p<0.05 vs. ctrl, #p<0.05 vs. pattern ($R^{hi}\varepsilon^{hi}D^{lo}$). † p<0.01 between marked groups. (D) Normalized collagen volume. ***p<0.001 vs. all other groups. (E) Collagen integrated density. *p<0.05 vs. all other groups. #p<0.05 between marked groups. N = 3-4 replicated gels per condition.

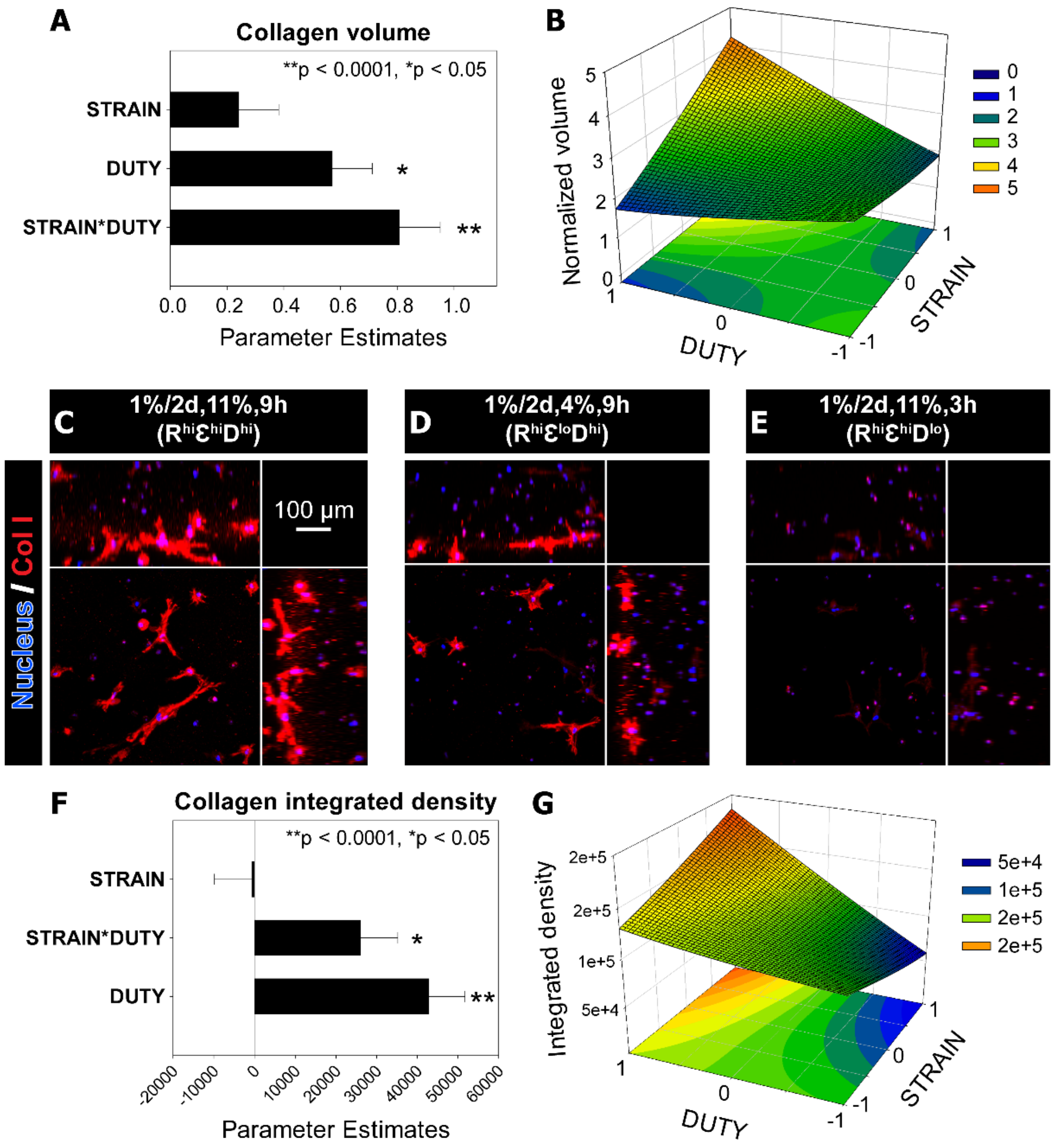


Figure S6. Parametric models of MSC responses of additional collagen metrics from two-week validation culture with RATE at 1%/2days. (A and F) Summaries of parameter estimates for (A) collagen volume (normalized to static ctrl) and (F) collagen integrated density. Refer to Table S4 for the prediction expressions for each response metric. Error bars represent standard error. N = 3-4 replicated gels per condition. $^{**}p < 0.0001$, $^{*}p < 0.05$. (B and G) The response surfaces of the STRAIN*DUTY interaction for (B) collagen volume and (G) collagen integrated density. 2D projection of the response surface is shown at the bottom of plot. (C-E) Representative maximum intensity projections of cells stained with collagen (Col I) for (C) Pattern ($R^{hi}Ɛ^{hi}D^{hi}$), (D) Pattern ($R^{hi}Ɛ^{lo}D^{hi}$), and (E) Pattern ($R^{hi}Ɛ^{hi}D^{lo}$). Peripheral images are side view projections.

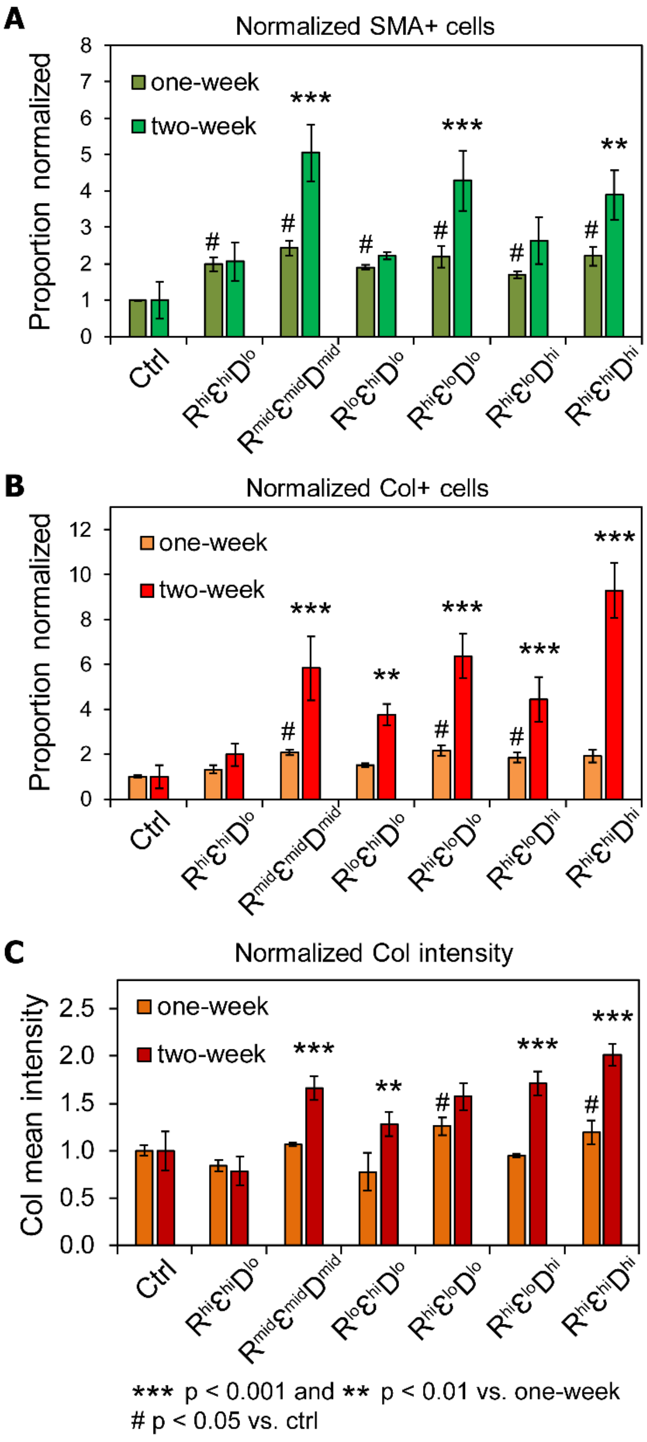


Figure S7. Comparison of MSC responses between the initial one-week screening and two-week validation culture. Results from each condition were normalized to that of the corresponding static control for valid comparison of (A) SMA+ proportion, (B) Col+ proportion, and (C) Collagen mean intensity. ***$p<0.001$ and **$p<0.01$ vs. corresponding one-week condition. #$p<0.05$ vs. ctrl. N = 3-4 replicated gels per condition.

Movie S1. MSC-laden PEG-NB hydrogels (left and right) under large deformation from the OSTE-PDMS membrane that is cyclically and pneumatically actuated, resulting in high tensile strain in the gel (i.e., 16% as shown in Figure 1C).
